# Itinerant lifestyle and congregation of lesser kestrels in West Africa

**DOI:** 10.1101/2022.08.12.503182

**Authors:** Lina Lopez-Ricaurte, Wouter M.G. Vansteelant, Jesús Hernández-Pliego, Daniel García-Silveira, Susana Casado, Fernando Garcés-Toledano, Juan Martínez-Dalmau, Alfredo Ortega, Beatriz Rodríguez-Moreno, Javier Bustamante

## Abstract

Migratory birds often spend a large proportion of their annual cycle in their non-breeding areas. While some species are sedentary after arriving in their non-breeding areas, others engage in itinerary or nomadic movements. Using GPS-tracking we aimed to investigate the little known non-breeding movements of the lesser kestrel *Falco namumanni* in sub-Saharan Africa. We segment non-breeding tracks (n = 78 tracks by 54 individuals) into staging events (131± 25 days), itinerant movements between staging sites (11 ± 10 days), and non-directed exploratory movements (6 ± 5 days). We then describe timing and directionality of itinerary movements by male and female kestrels, and explore shifts in habitat use throughout the non-breeding season. Regardless of sex, lesser kestrels spent on average 89% of the non-breeding season at 2-3 staging sites in West Africa. Upon arrival at the end of September, kestrels used disparate sites throughout the Sahel. By December, however, they congregated into two clearly delineated areas in Senegal and along the Malian-Mauritanian border. The birds stayed longer and showed greater daily activity in the latter areas, which were situated closer to wetlands compared to their first and intermediate ones. While 20 of 24 individuals tracked across multiple annual cycles showed consistent staging sites, a few switched between the Senegal and Mauritanian/Malian staging clusters. These results show that rather than tracking suitable climatic conditions southward, lesser kestrels survive the West African dry season by moving into wetland areas within the Sahelian climatic belt. Our findings match reports of lesser kestrel super-roosts in West Africa and stress the conservation value of wetlands along the Senegal River and the Inner Niger Delta in West Africa for the Spanish lesser kestrel population. These areas host 81% of lesser kestrels during 20% of their annual cycle.

## Introduction

The ecology of migratory animals is shaped by conditions in more than one part of the world: their breeding grounds, non-breeding quarters and migratory routes (Bildstein, 2006; Schofield et al., 2010; Cherry et al., 2016). Owing to their reliance on multiple areas across their annual cycle, they are in ‘multiple jeopardy’ (Sanderson et al., 2006; Gilroy et al., 2016). Importantly, long-distance migrants may spend the majority of their annual cycle in their non-breeding areas (Newton, 2004; Studds & Marra, 2005; Salewski & Jones, 2006), and conditions in those areas can have carry-over effects on survival and reproduction (Marra et al., 1998; Newton, 2004; Norris et al., 2004). Despite this, many aspects of migrants’ non-breeding ecology, in particular birds moving into tropical latitudes, and the potential threats they face in such distant, often hard to access non-breeding areas, remain to be learned (reviewed in Wilcove & Wikelski, 2008; Stanley et al., 2021).

Several billion land birds of about 200 species from the Palearctic spend the boreal winter in sub-Saharan Africa (Dean, 2004; Salewski & Jones, 2006). It has been suggested that an increased mobility during non-breeding is needed for birds to track seasonally shifting resources across large areas (Sinclair, 1978; Lack, 1983; Salewski et al., 2002). In addition, and contrary to the breeding period, birds have more freedom to move during the non-breeding season, when they are not tied to a nest site. This freedom to move historically hampered our ability to study such species for an extended period (Bildstein, 2006). Nowadays, however, advanced tracking technology allows the study of a wide range of Palearctic-African migrants, including species with highly mobile lifestyles (Smith et al., 2011; Vickery et al., 2014; Norevik et al., 2019). Furthermore, migrant birds have been differently affected by environmental changes in sub-Saharan Africa, depending on non-breeding distribution (Sanderson et al., 2006; Zwarts et al., 2009; Vickery et al., 2014). As such, unravelling non-breeding movements and space use is fundamental to flyway conservation.

Bio-logging has confirmed different movement strategies and great interspecific behavioural flexibility during the non-breeding period (Bildstein, 2006; Salewski & Jones, 2006; Norevik et al., 2019). Some species, like pied flycatchers *Ficedula hypoleuca* (Ouwehand et al., 2016) and common redstarts *Phoenicurus phoenicurus* (Kristensen et al., 2013), remain on a single territory (often covering smaller areas than their breeding home ranges) during the whole non-breeding period, and remain faithful to these areas between years, a strategy known as non-breeding residency (Newton, 2008). Conversely, some species show high non-breeding mobility within or between years, performing extensive movements over a large non-breeding area and even visiting different destinations year after year (Dean, 2004; Newton, 2008; van Wijk et al., 2016). A nomadic lifestyle may be prevalent in semi-arid heterogeneous environments (e.g. deserts) where resources are patchy in space and time and determined by inter-annual fluctuations in rainfall (Jensen, 1972; Dean, 2004).

An intermediate strategy to residency and nomadic lifestyles is itinerancy, occupying two or more residence areas during the course of the non-breeding season in succession and consistently staging in the same general areas over consecutive years (Moreau, 1952, 1972). Using the same residency sites each year, such migrants may cope with inter-annual variation in resource availability by adjusting the time they spend in each of these areas. For example, as aridity increases in areas north of the equator as the season progresses, birds may follow the seasonal shifts of food abundance southwards to more benign habitats (Trierweiler et al., 2013; Thorup et al., 2017). This behaviour has been reported in the Palearctic-African bird migration systems (e.g. reed warblers *Acrocephalus arundinaceus*, Lemke et al., 2013; lesser spotted eagles *Clanga pomarina*, Meyburg et al., 2015; willow warblers *Phylloscopus trochilus*, Lerche-Jørgensen et al., 2017; pallid swifts *Apus pallidus*, Norevik et al., 2019; Montagu’s harriers *Circus pygargus*, Trierweiler et al., 2013; Schlaich, 2019).

There seems to be some interindividual variation in itinerant species, where some individuals behave as winter residents and others as itinerants. For example, 3 individual turtle doves *Streptopelia turtur* used one residency site, while 2 were itinerant (Eraud et al., 2013). The same pattern was also shown by 1 out of 6 tawny pipits *Anthus campestris* (Briedis et al., 2016), by 4 out 129 Montagu’s harriers *C. pygargus* (Schlaich, 2019) and by 2 out of 10 marsh harriers *Circus aeruginosus* (Vansteelant et al., 2020). Furthermore, age and sex differences in non-breeding schedules and habitat use have been reported. For example, female American kestrels *Falco sparverius* arrive first in South Florida’s non-breeding territories and occupy better habitats than males (Smallwood, 1988; Ardia & Bildstein, 1997). Evidence from ringing indicates that males yellow wagtails *Motacilla flava* winter further north than females (Wood, 1992). Yet, assessing individual and sex differences in non-breeding strategies is often challenging due to the technological challenge of tracking multiple individuals of both sexes across consecutive non-breeding periods.

In this study, we use an extensive GPS tracking dataset to investigate non-breeding mobility and space use in the lesser kestrel; a small trans-Saharan migratory raptor that breeds in colonies from southern Europe and northern Africa to China. During the boreal winter, it has an Afrotropical distribution (Ferguson-Lees & Christie, 2001), although some populations in the Mediterranean exhibit partial migration, with a minority of individuals -mostly males- being non-migratory (Negro et al., 1991). Previous studies using geolocators (Rodríguez et al., 2009; Catry et al., 2011) and satellite telemetry (Limiñana et al., 2012) identified non-breeding areas of lesser kestrels from the western breeding range. Moreover, Catry et al. showed interindividual variation in non-breeding movements with one female using one residency site while three others used two or more sites (Catry et al., 2011). Recent GPS-tracking studies have revealed that lesser kestrels from Mediterranean populations migrate in a broad front across ecological barriers (e.g., the Mediterranean Sea and Sahara Desert) and that there is strong connectivity between breeding and non-breeding areas (Sarà et al., 2019; López-Ricaurte et al., 2021). Furthermore, during the non-breeding season, lesser kestrels are known to aggregate in super roost wetlands within western Senegal (e.g. Kaolac and Khelkom) were several tens of thousands of individuals have been observed in late non-breeding period (mid-January) (Pilard et al., 2011; Augiron et al., 2015). Analysis of pellets collected at such super roosts revealed that lesser kestrels feed almost exclusively on insects (mostly Orthoptera) in these areas. Field observations suggest they hunt in flocks over arable land, shrub savannah and grassy savannah (Pilard et al., 2011; Augiron et al., 2015). However, we still know relatively little about lesser kestrel’s non-breeding ecology outside these areas and whether they also form such super roosts elsewhere.

This study aims to describe the non-breeding movements of lesser kestrels from the Spanish breeding population in West Africa, and to test for potential differences in the mobility and timing of intra-African movements between sexes. Because of the high movement ability of these flight generalist species (Lopez-Ricaurte et al., 2021; Bildstein, 2017), we expect lesser kestrels to show itinerant movements within West Africa, moving progressively further south over the season to follow the seasonal shifts in insect abundance (c.f., other locust-eating steppe birds, Berthold 2002, 2004; Trierweiler et al., 2013; Schlaich et al., 2016). Finally, we characterise land use at their staging sites using the GlobCover land use map (2009) (resolution 300m) in a similar way to previous studies (Trierweiler et al., 2013; Schlaich, 2019).

## Materials and Methods

### Tagging and tracking

Lesser kestrels were tagged using two models of solar GPS-UHF biologgers (GPSminiDatalogger, Microsensory LS, Córdoba, Spain; and NanoFix GEO+RF, Pathtrack Ltd., Leeds, UK.). Birds were tagged at 20 breeding sites across Spain by different organisations (GREFA; The Spanish Society of Ornithology, SEO/BirdLife; Terra Naturalis, and Doñana Biological Station, EBD). The GPS-UHF biologgers weighing ca. 5.5 g (including harness, ~3.8 % of weight at capture, males = 146.0 g ± 35 SD; females = 148.0 g ± 29 SD) were attached as backpacks with a Teflon harness. As GPS-UHF biologgers were deployed for different projects by different teams, they were programmed with different schedules (see Supplemental methods for details). Locations were stored on-board and later downloaded via a UHF base station placed near the breeding colony.

We relied on 78 non-breeding tracks of 54 adult birds (25 males and 29 females) (Supplemental Figure 1). Twenty three individuals provided tracks for two consecutive non-breeding seasons and 1 individual for 3 non-breeding seasons.

### Annotating non-breeding movements

All data were resampled to a 1-h interval, allowing deviations up to 20 min. By resampling, we also avoided bias in our calculations of movement parameters due to the variability in sampling rates (Shamoun-Baranes et al., 2017). After resampling, we analysed 167,793 hourly segments, from which 108,792 were recorded by day and 59,001 by night. We determined arrival to and departure from the non-breeding grounds based on daily movement metrics (see López Ricaurte et al., 2021 for full details). We studied non-breeding movements within Africa by interpreting the tracks using QGIS (cf. Trierweiler et al., 2010; Schlaich, 2019). We annotated as (1) ‘resident days’, each day in a group of ≥ 3 days in which the bird stayed stationary at a site (using roosts less than 10 km apart on consecutive nights), (2) “transit days”, when the individual performed a directional flight away from a site without returning, and (3) “exploratory days”, when birds performed non-directed movements away from a site but returned to a previous roost (could last one or several days) (Supplemental Figure 2).

For mapping, we determined the centroid of each site where the bird was resident as the median latitude and longitude of all positions at this site (hereafter staging sites). The staging sites were subdivided into three categories: the ‘first’ staging site south of 17° N used upon arrival from the post-breeding migration, ‘last’ as the last staging site used before the onset of the pre-breeding migration, and ‘intermediate’ all other consecutive sites used between first and last staging sites (could be one or several). Sometimes individuals remained highly mobile for 1 – 2 days upon arriving to West Africa and before moving into their first staging site (5 individuals). Such movements were classed as ‘exploratory days’. The arrival date to the non-breeding grounds in such birds was determined as the date on which the individual moved into the first staging site. Three individuals stayed the whole non-breeding period at a single staging site; and these were classified as last staging sites (because birds using more than one site tended to spend the most time at the last site).

### Non-breeding schedules and movement metrics

We computed arrival and departure dates to and from West Africa as the first and last day that the bird moved into the first and last staging sites, respectively, the total duration within West Africa (in days), the total number of resident, transit and exploratory days per individual and the daily distance covered (mean of the sum of successive distances between the first and the last GPS fix of a day). For each first, intermediate, and last staging site we quantified the (1) arrival and departure dates to each staging site (2) the staging duration per site (3) mean cumulative daily distance, (4) mean trajectory ground speeds from each GPS fix to the previous (i.e., the speed between consecutive fixes), (5) the daily time spent flying/sitting and (6) the daily proportion of daylight period spent flying/sitting. To calculate (5) and (6) we determined ‘flying’ segments as hourly segments with a ground speed ≥ 5 km/h.

### Statistical analyses

We tested for sex differences in arrival, departure dates, and duration of the stay in West Africa using Linear Mixed Models (LMMs), allowing for random intercepts per bird. Visual inspection of residual plots indicated that a Gaussian error distribution and identity link function provided the best model fit.

Differences in the daily distance covered among types of days (resident, transit, exploratory) and between sexes were investigated using Generalised Linear Mixed-effect Models (GLMMs). We also tested for differences in timing of itinerary movements (arrival and departure dates) and mobility (daily distance, duration of the stay and % of time spent flying) between consecutive staging sites and sexes using GLMMs. The percentage of time spent flying was arcsine-square-root transformed to get a proper fitting of the models. We added the variable ‘staging site’ (with three levels: first, intermediate, last) and ‘sex’ as fixed factors, and bird identity and non-breeding cycle (i.e. 2018-2019,2019-2020,2020-2021) as random effects in all models. For pairwise comparisons, we used Tukey’s HSD (honestly significant differences) tests, whereby we considered an effect to be statistically significant if p ≤ 0.05.

### Habitat composition and use

We estimated the overall non-breeding home range for the Spanish lesser kestrel population as the 100% minimum convex polygon (MCP) across the centroid of all staging sites. We excluded one site that was situated far south of all kestrel staging sites in Guinea, resulting in a final sample of 196 staging sites. To examine lesser kestrels’ broad habitat use across West Africa, we used the GlobCover 2009 V2.3 land use map at 300 m resolution (Defourny et al., 2009). First, we calculated the percentage of available land cover types within the MCP non-breeding area. Then, the habitat types used by lesser kestrels were determined by projecting 1-h GPS daylight fixes onto the GlobCover map and extracting habitat type for each hourly location. Transit and exploratory days were excluded from the habitat use analyses.

## Results

### General description of non-breeding movements

The average arrival date to West Africa was September 30 ± 12 days, and the average departure date was February 24 ± 14 days (Table 1). The number of days spent in the non-breeding range was 147 ± 17, corresponding to 40% of the total annual cycle (Supplemental Table 1). We did not find any significant sex differences in arrival, departure dates and duration of the stay in West Africa (Table 2). From the time spent in Africa, 131 ± 25 days were resident (sedentary at a staging site, 89% of the non-breeding period), 11 ± 10 days were transit (itinerant movements between staging sites, 7% of the non-breeding period), and 6 ± 5 days were exploratory (4% of the non-breeding period). The mean daily distance covered during resident days was 40.41 ± 36.60 km, during transit days was 80.89 ±71.01 km, and that of exploratory days was 98.03 ± 73.96 km, and these differences were significant (GLMM: *F* = 7825.90, P ≤ 0.001; Fig. 1A). Mean daily distance across all days did not differ significantly between males and females (GLMM: *F* = 1.31, *P* = 0.25).

**Figure 1.**
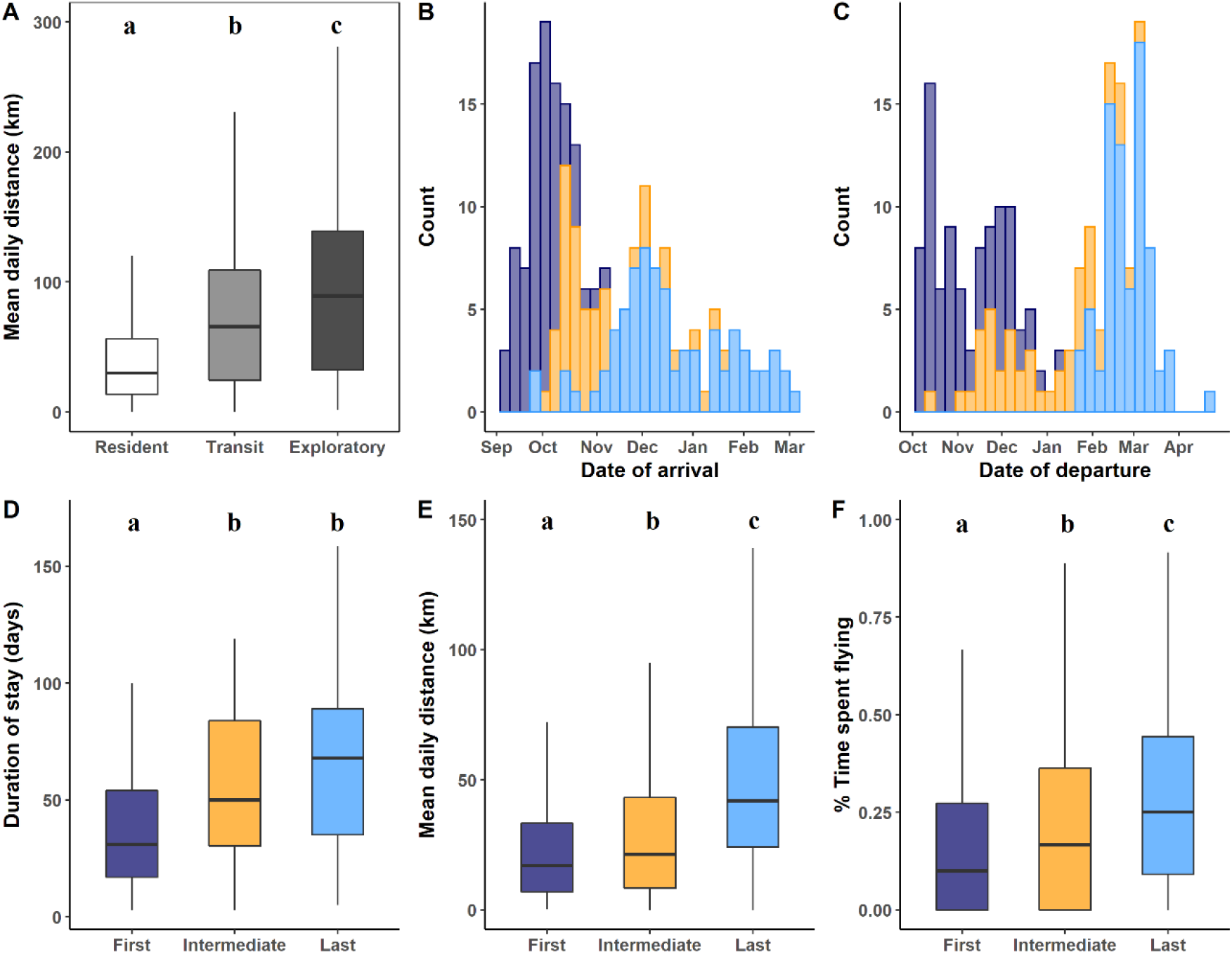
**(A)** Mean daily cumulative distance during resident, transit and exploratory days, **(B)** arrival dates, **(C)** departure dates, **(D)** total days spent per staging site type, **(E)** Mean daily distance covered per site, and **(F)** percentage of daylight hours with flight activity (segments with ground speed ≥ 5 km/h). Colours indicate the type of staging site: first sites in purple, intermediate sites in yellow, and last sites in blue. The letters above represent significant differences by Tukey HSD post-hoc tests at the 0.05 significance level. Groups sharing the same letter are not significantly different.

**Table 1.** Arrival and departure dates and duration of lesser kestrels at the non-breeding grounds.

| Variable | N<br>(tracks/individuals) | Mean (± SD) | Min | Max | Range |
| --- | --- | --- | --- | --- | --- |
| Arrival | 78/54 | Sep 30 ± 12 days | Sep 9 | Nov 9 | 61 |
| Departure | 78/54 | Feb 24 ± 14 days | Jan 24 | Apr 18 | 100 |
| Duration (days) | 78/54 | 147 ± 17 days | 92 | 200 | 108 |

**Table 2.** Linear mixed models to test for sex differences in arrival date, departure date and duration. Individual identity (ID) was included as a random factor in the models.

| Response variable | Fixed effects | Estimate | SE | t | P |
| --- | --- | --- | --- | --- | --- |
| Arrival date | Intercept | 272.14 | 2.03 | 133.63 | $\leq 0.001$ |
|  | Sex (male) | 2.73 | 2.86 | 0.95 | 0.34 |
| Departure date | Intercept | 56.73 | 2.61 | 21.72 | $\leq 0.001$ |
|  | Sex (male) | -2.33 | 3.74 | -0.62 | 0.53 |
| Duration (days) | Intercept | 149.18 | 3.26 | 45.67 | $\leq 0.001$ |
|  | Sex (male) | -4.75 | 4.58 | -1.03 | 0.30 |

*N = 78 observations, ID = 54 individuals*

### Temporal and movements patterns at staging sites

The staging sites of tracked birds were located across West Africa (Senegal, Mauritania and western Mali) between ca. 13.5° and 17.5° N and −16° and −4° W (Fig. 2A). We observed either a westward or an eastward individual movement pattern from the first to last staging site as the non-breeding period progressed (Fig. 2B, 2C). First staging sites were mainly distributed at central longitudes, intermediate sites were more spread out, and the last sites fell in two distinct eastern and western clusters. On average, individual kestrels used 2.5 ± 0.71 staging sites, ranging from 1 (3 of 78 tracks, 3.84%) to 4 (5 of 78 tracks, 6.41%) (Fig. 2D). The median arrival date at the first sites was September 29, at the intermediate sites October 20, and at the last sites November 20 (Fig. 1B). The number of staging sites used did not differ significantly between sexes (GLMM: *F* = 1.05, *P* = 0.31; Fig. 3A). The median departure date from the first site was October 26, from the intermediate site November 1, and from the last site February 23 (Fig. 1C) with no differences between sexes (GLMM: *F* = 0.96, *P* = 0.33; Fig. 3B).

**Figure 2.**
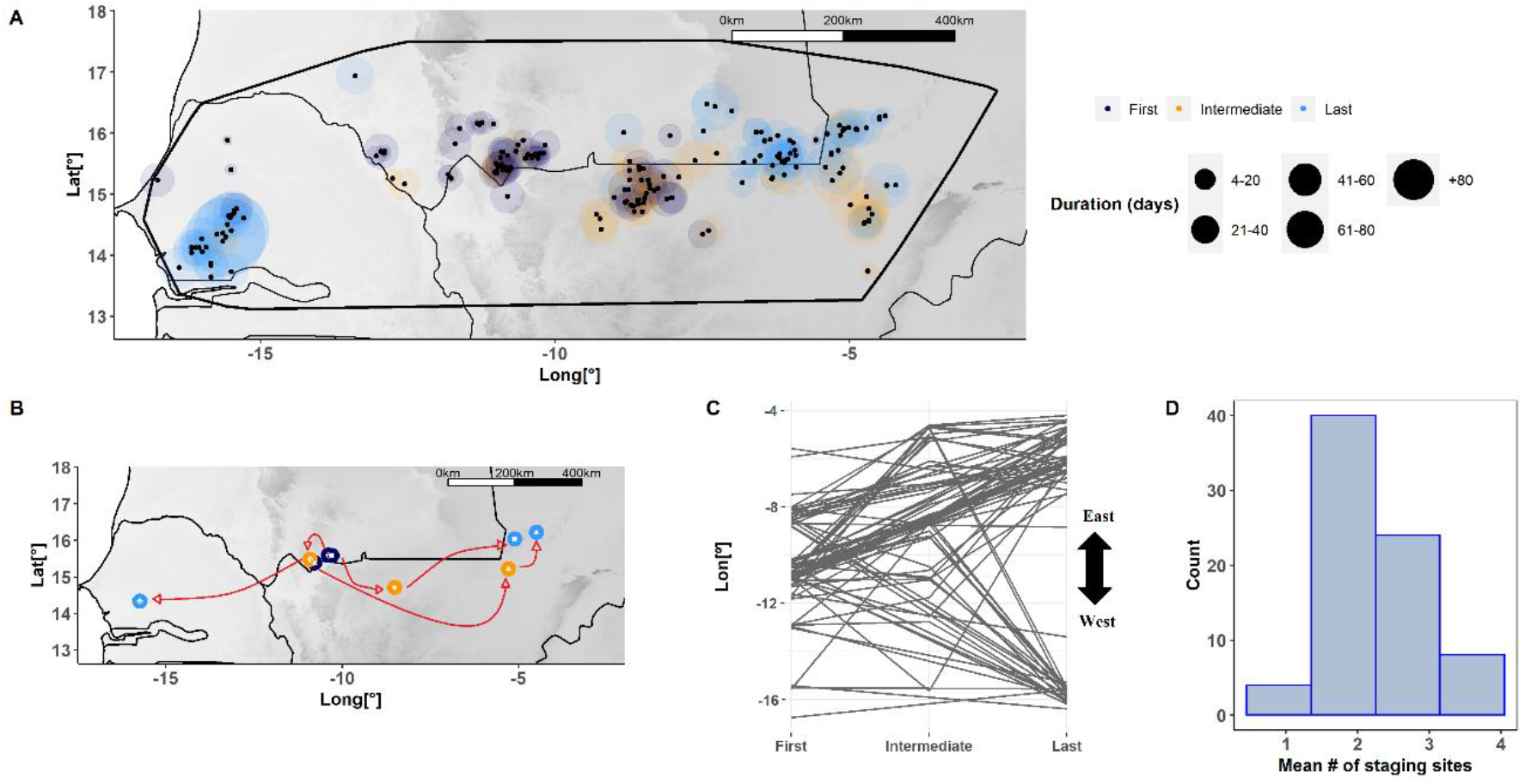
**(A)** Map showing the location of lesser kestrel staging sites in West Africa. Colours indicate the type of staging site: first sites in purple, intermediate sites in yellow, and last sites in blue. The black dot represents the centroid of each site (i.e. median latitude and longitude of all positions at a site). The 100% MCP representing the tracked kestrels’ non-breeding range is also depicted. **(B)** Example of three individual tracks representing the typical movements of lesser kestrels during the 2018-2019 non-breeding period, i.e. birds arriving within central West Africa and then dispersing into the west or east staging sites. Large circles represent staging sites (first in purple, intermediate in yellow, last in blue), and symbols represent different individuals: female 4178696_RVFN (diamond), female 4170460_4JT (square) and male 4173803_RJA1 (triangle). Red arrows connect subsequent staging sites of the same individual. **(C)** First, intermediate and last staging sites used by each individual related to their mean longitude. **(D)** Histogram of the mean number of staging sites used by each individual.

**Figure 3.**
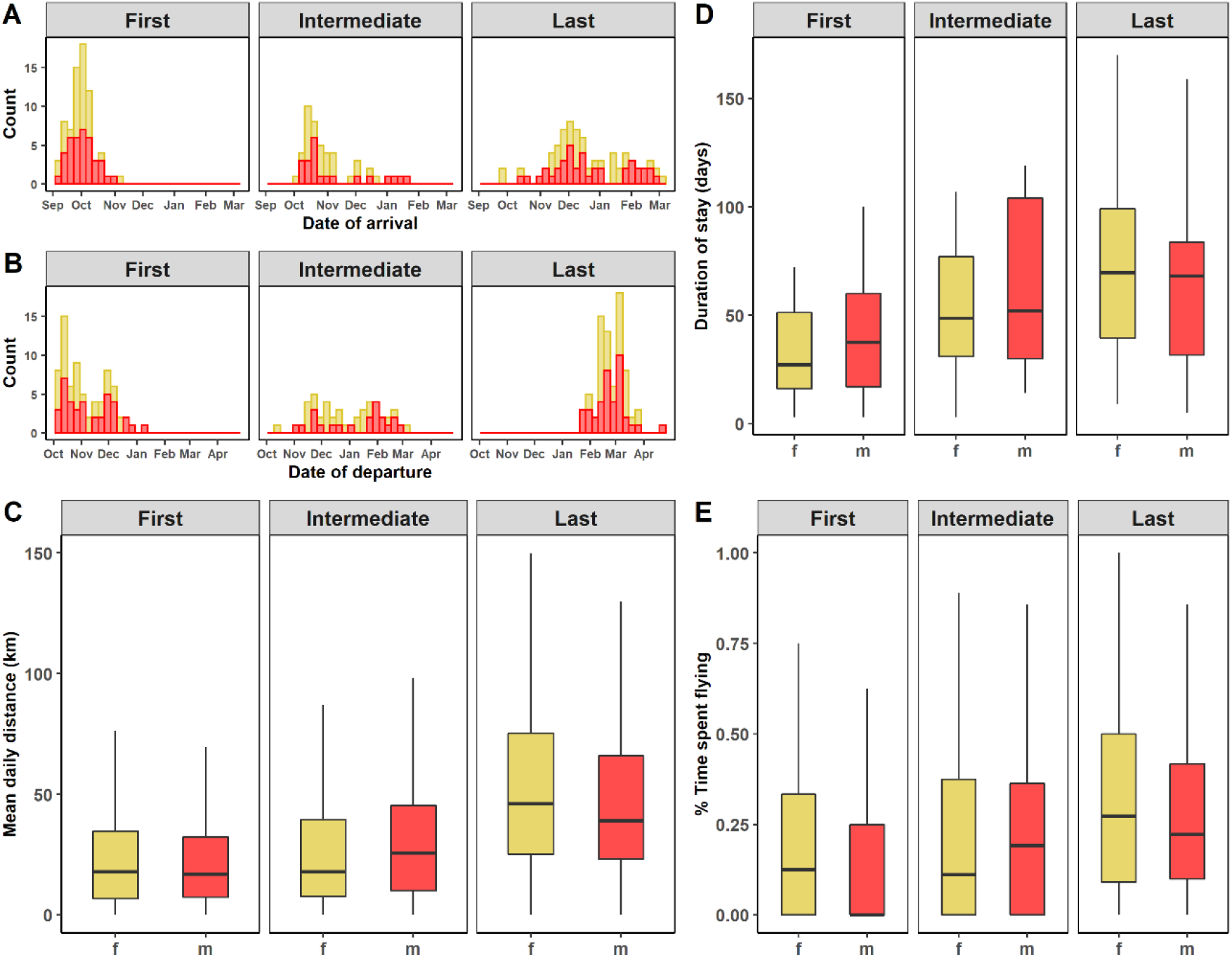
**(A)** Histogram of arrival dates of lesser kestrel males and females at first, intermediate and last staging sites, (females in yellow, males in red). **(B)** Histogram of departure dates. **(C)** Histogram of mean daily distance per site. **(D)** Mean duration of stay according to sex at first, intermediate and last staging sites, and **(E)** Mean percentage of daylight hours with flight activity (segments with ground speed ≥ 5 km/h) in relation to sex.

The mean duration of the stay was significantly different among consecutive staging sites (GLMM: F 41.38, P < 0.001). Kestrels spent on average 36 ± 21 days on the first staging site, 52 ± 32 days at intermediate sites, 68 ± 36 days at their last staging site, corresponding to 10%, 14% and 20% of the total annual cycle, respectively (Supplemental Table 1), with no differences between sexes (GLMM: F=0.007, P = 0.93; Fig. 3D). Birds spent significantly longer time at intermediate and last sites than at first sites (Fig. 1D). The daily distance covered and the percentage of time spent flying were significantly different between sites (mean daily distance GLMM: F=10927.77, P < 0.001; percentage of time spent flying: GLMM =9720.97, P < 0.001). Birds covered significantly longer daily distances and spent more time flying at the last site relative to the first and intermediate sites (Fig. 1E and F) with no sex differences (mean daily distance GLMM: F=0.22, P = 0.63; percentage of time spent flying GLMM: F=0.94, P = 0.33) (Fig. 3C, E).

At the end of the non-breeding period, lesser kestrels typically congregated in two clearly delineated clusters, one in Senegal, on the western side of the non-breeding range, and another on the Mauritania-Mali border eastern side. Out of 78 tracks, 32% converged close to Kaolack, Khelkom and Prokhane (west Senegal), including 26% of all females and 35% of all males. In addition, 49% congregated in Bassiknou and Djiguenni (Mauritania), including 59% of all females, and 44 % of all males. Furthermore, 19% converged in Lere and Mopti (Mali), including 15% of all females and 21% of all males (Fig. 4).

**Figure 4.**
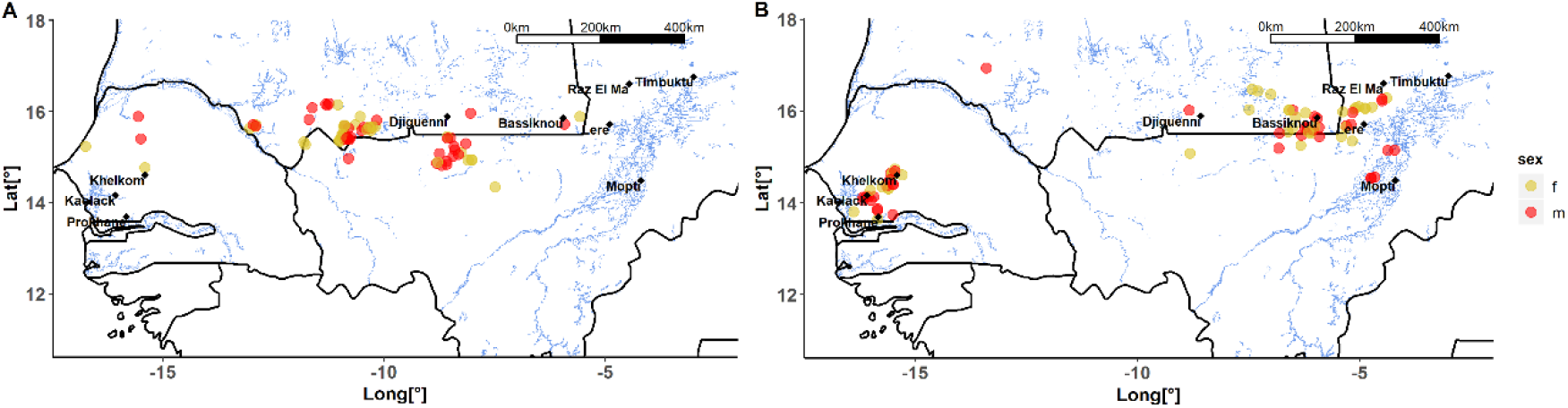
Map showing (**A**) first and (**B**) last staging sites according to sex (females yellow and males in red dots). Blue coloured lines indicate the inland water present in Senegal, Gambia, Mauritania and Mali. Black diamonds indicate the region closer to lesser kestrels staging sites. In central Senegal: Khelkom (also known as Mbégué), Kaolack and Prokhane (near the border with Gambia); in southwestern Mauritania: Bassikounou and Djiguenni in the region of Hodh Ech Chargui, and in northeastern Mali: Mopti, Timbuktu, Raz El Ma and Lere. A notable cluster of sites closer to floodplains (Inner Niger Delta), wetlands in eastern Mauritania, and coastal wetlands, notably Sine Saloum in Senegal and the Gambia, can be observed at the end of the non-breeding period.

Of the birds tracked during two consecutive years (n=23) and three consecutive years (n=1), (n= 8 females and 16 males), 33% (8 out of 24 tracks) consistently revisited the same last site (maximum distance between sites < 20 km) in consecutive years. In addition, 50% (12 out of 24 tracks) of lesser kestrels used sites near the one visited in an earlier non-breeding period (< 200 km apart). Most of these birds (83%, 20 out of 24 tracks) used the same general non-breeding area in consecutive non-breeding cycles (Senegal vs Mauritania-Mali border). Finally, 16% (4 out of 24 tracks, all adult males) changed the last staging site (> 300 km apart) (Fig. 5).

**Figure 5.**
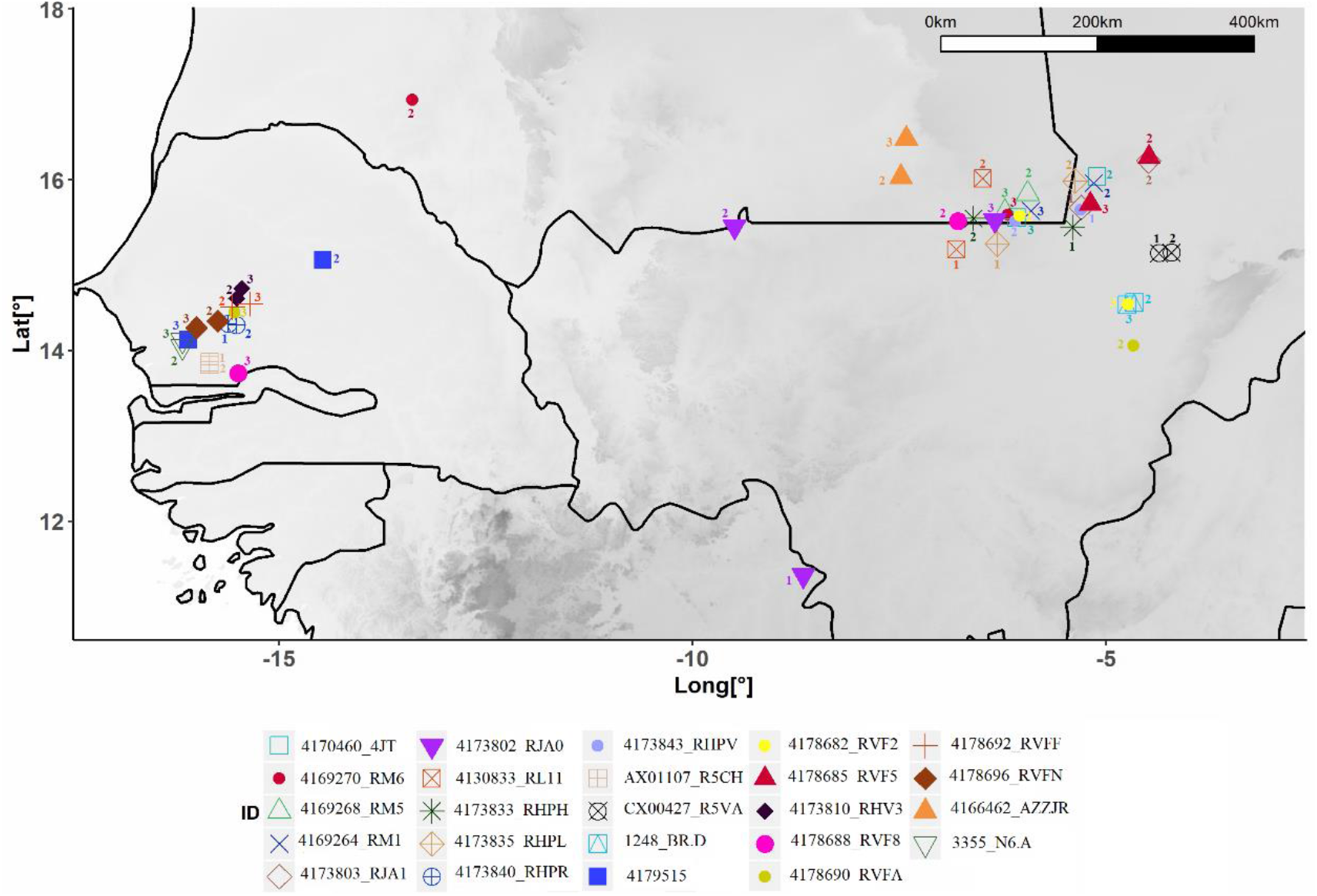
Last staging sites in West Africa of 24 adult lesser kestrels (represented with different colours and symbols) tracked during consecutive years. 83 % of these kestrels consistently spent the last period of their stay in West Africa in the same or near the same area as the previous year. Numbers correspond to the year cycle: 1 to 2017-2018, 2 to 2018-2019 and 3 to 2019-2020. Bird 4169270_RM6 (solid circles, dark red) moved from western Mauritania in the first non-breeding period to the Mauritania-Mali border in the second non-breeding period; bird 4173802_RJA0 (solid inverted triangle, purple) moved from Guinea in the first non-breeding period to western Mauritania in the second non-breeding period and to the Mauritania-Mali border in the third non-breeding period; bird 4178688_RVF8 (solid circles, pink) moved from the Mauritania-Mali border in the first non-breeding period to Gambia in the second non-breeding period, and bird 4178690_RVFA (solid circles, kaki) moved from Mali in the first non-breeding period to Senegal in the second non-breeding period.

### Habitat use

Fourteen out of 23 GlobCov categories were available across the potential non-breeding range of lesser kestrels. Grassland, mosaics of vegetation (veg/crop), mosaics of cropland (crop/veg) and crops were the most frequent categories in the region (23%, 21%, 16%, 10%, respectively) (a description of the GlobCover categories is included in Supplemental Table 2).

Our results for habitat suggested some shifts in habitat used along successive staging sites. In the first sites, 11 GlobCov categories were used. We found that relative to the habitats available in West Africa, kestrels used veg/crops, crops/veg, crops, sparse vegetation, followed by mosaics of shrubland (shrub/grass) and grassland. The least used habitat types relative to what is available were water, bare, shrub, mosaics of grassland (grass/shrub) and forest. 12 GlobCov categories were used at intermediate, and 14 categories at last sites. At intermediate sites, they primarily used veg/crops and crops/veg, followed by crops, shrub/grass, grassland and sparse. At last sites, they primarily used veg/crops, crops/veg, shrub/grass and also, grassland, sparse and bare habitats. We observed that the use of mosaics of vegetation (veg/crops), mosaics of cropland (crops/veg) and crops decreased gradually from the first to last sites. Conversely, the proportion of time spent by kestrels in bare, urban and moister habitats (water, wet soil and mangrove) increased from first to last sites (Supplemental Figure 3).

## Discussion

We found that lesser kestrels have an itinerant lifestyle during the non-breeding period in West Africa. This is an intermediate strategy between residency at a single site and a nomadic lifestyle. Lesser kestrels used a small number of staging sites (two or three) in one non-breeding season, to which they showed fidelity in consecutive years, particularly to the last staging sites. Kestrels spent most of their non-breeding time (89%) in those sites. Conversely, they spent a relatively small proportion of time moving between sites (7%) or engaging in seemingly exploratory movements (4%). Lesser kestrels arrived to their first non-breeding staging sites at the end of September and they spent 147 days in West Africa before departing north, typically in the last week of February. Moreover, males and females followed a similar non-breeding strategy, with no significant differences in the timing of itinerary movements or movement metrics at successive staging sites. We also observed that kestrels spread out over the interior of West Africa at arrival – from the northeast Senegalese border to the easternmost part of the Mauritania-Mali border. Contrary to our expectations and to other locust-eating steppe birds, however, kestrels did not move along a north-south axis in West Africa (Trierweiler et al., 2013; Schlaich et al., 2016). Instead, they moved either westward or eastward within the Sahel region, often using 1-2 intermediate sites before converging at staging sites in inland wetlands in Senegal or in eastern Mauritania at the border with Mali, respectively (Fig. 1 A, B, C; Fig 4B).

The minimum number of days used to define a site as a staging site influences the final number of staging sites per individual. Our classification of staging sites is reliable because we found that repeatedly tracked birds were faithful to sites identified based on the ‘3-day rule’. Upon arrival, lesser kestrels typically settled for 1 month just south of the Sahara Desert. We suspect that after traversing ecological barriers with limited possibilities for fuelling during the post-breeding migration, birds may stop at the first suitable place they encounter in the northern Sahel at the end of the rainy season (Trierweiler et al., 2010; Vansteelant et al., 2020). Lesser kestrels may then remain there until the region becomes too inhospitable due to increasing aridity (Salewski et al., 2002). During the transition from the wet to the dry season, kestrels move out of the most arid regions, moving either westward or eastward into areas with more abundant and larger water bodies within the Sahel (e.g. coastal wetlands, Inner Niger delta, wetlands in eastern Mauritania), which dry out later and may suppress the adverse effects of the dry season for several months (Studds & Marra, 2005). Such least arid habitats may sustain vegetation and associated insect life for longer in the dry season, enabling kestrels to spend a large part of their non-breeding period at these sites (Zwartz et al., 2009; Vafidis et al., 2014). While the kestrels’ longitudinal itinerary movements differ from the predominantly latitudinal movements of other locust-specialists and aerial hunters (swifts, Akesson et al., 2012; swallows, Norevik et al., 2019, harriers, Trierweiler et al., 2013; Schlaich et al., 2016), east-west movements within the Sahel have also been reported for other species (e.g. tawny pipits *A. campestris*, Briedis et al., 2016; turtle doves *S. turtur*, Eraud et al., 2013).

Lesser kestrels exhibited little individual variation in non-breeding strategy with no sex differences in schedules or movement metrics. We found that only 3 tracks out of 78 (3.84 %) showed a strategy of non-breeding residency, staying the whole non-breeding period at the same staging site, while the great majority were itinerant. Lesser kestrels are food specialists (eating namely flying insects, e.g. grasshoppers during the non-breeding period, Pilard et al., 2011). Therefore they may rely on an itinerant strategy to track the fluxes of their prey, in contrast to food generalists who could feed on a diversity of prey items available in the same site (Salewski et al., 2002). Furthermore, we did not find any sex differences related to schedules or movement metrics. Sex-related differences in non-breeding schedules, movements and habitat use have been reported, particularly in species with size dimorphism. Males ruffs (*Philomachus pugnax*), the larger sex, winter further to the north than females, and their migration phenology is more advanced (van Rhijn 1991). Based on field observations, males wintering in the Senegal Delta started to fatten before embarking on northward migration three weeks earlier than females (Zwarts et al., 2009). Field-based studies on Marsh harriers wintering in the Inner Niger Delta showed that females use wetlands, likely due to their ability to hunt larger waterbird prey, while males focus more on small prey such as small mammals and grasshoppers in dryer areas Bijlsma et al., 2001). The lesser kestrel has a reversed size dimorphism, with females being 15% heavier than males (Cramp, S. & Simmons, 1980). However, such a size difference may not be enough to give females access to alternative resources than males, so they respond similarly to conditions in the Sahel.

Our tracked lesser kestrels spent the longest time (two months on average), covered longer mean daily distances, and spent more daylight hours flying in the last staging sites than at first and intermediate ones. We speculate three non-exclusive explanations: (1) the necessity of depositing fat during the last weeks prior to the return migration (Newton, 2008). (2) The high intraspecific competition derived from the massive individual aggregation in the last staging sites may imply an increased foraging effort. And (3) kestrels might use larger prey species at the last staging sites compared to the previous sites (e.g. grasshoppers species such as *Ornithacris cavroisi* or the larger desert locusts *Schistocerca gregaria* females, Mullié, 2021), requiring more time on the wing (more time flying vs. perch hunting smaller prey at first sites).

Lesser kestrels are insectivorous steppe birds using open grasslands and croplands at their breeding areas in the Mediterranean (Franco et al., 2004; Ursúa et al., 2005). Our results indicate that during the non-breeding season they use mosaics of grassy vegetation, shrubland, cropland, and sparse vegetation. Additionally, our results suggested some shifts in habitat use over the non-breeding period. Lesser Kestrels first stopped in crops and mosaics of habitat types along the Mauritania and Mali border. The most important non-breeding habitats at intermediate and last sites were mosaic habitat types (e.g. vegetation, cropland, shrubland, for more detail on these categories, see Supplemental Table 2) and more natural habitats such as grassland and sparse vegetation. This is in accordance with previous studies, which suggested that Palearctic migrants typically used heterogeneous rather than homogenous landscapes in their wintering grounds (Salewski & Jones, 2006). For example, Khelkom (Senegal) was protected under Senegalese law until 1991. Since then, the area was gradually cleared as rangeland and for groundnut development until 2004. The resulting mosaic of cropped areas, arable land and regenerating natural savannah has become an ideal habitat for grasshoppers. In turn, the area has attracted massive numbers of insectivorous bird species such as white storks *Ciconia ciconia* (3.500 individuals; representing 1.75% of the flyway population), Montagu’s harriers *C. pygargus* (5.000-6000; 16%) and lesser kestrels *F. naummani* (5.000; 10%) (Mullié & Guèye, 2010).

There are potential caveats to the interpretation of the habitat use data in the study. It should be noted that the map consists only of 22 classes and that the study region features numerous other habitat types that are likely to be important for lesser kestrels because large concentrations of locusts may occur there (e.g. laterite plateaus, estuaries, Schlaich, 2019). Therefore, it is likely that our interpretation of habitat use may underestimate the importance of other habitats used by lesser kestrels, and in any case, it is hard to translate land use to ecologically relevant parameters such as prey availability. We are confident that some patterns revealed from GlobCover data are robust, such the apparent avoidance of widely available grassland at the start of non-breeding period and the increasing diversity of habitat use from first to last staging sites. However, further investigation of habitat preferences and land cover change would require other more detailed habitat maps to match better relevant resources for kestrels (e.g. insect distribution, fires) in West Africa and, ideally, ground-truth such products in the field to improve accuracy (Vickery et al., 2014).

### Conservation implication

In the late 1960s and early 1970s, numerous Afro-Palearctic migrants –including the lesser kestrel – showed a sharp population decline (Sanderson et al., 2006; Iñigo & Barov, 2010). During the first decade of the 21st century, there have been several years of higher rainfall in the Sahel which may have led to population recovery of Afro-Palearctic migrants (Nevoux et al., 2008), including probably the lesser kestrel. However, this positive trend has not been maintained in subsequent years, and like numerous steppe and grassland birds, the lesser kestrel is now declining in Spain (Bustamante et al., 2020). While land-use change in the breeding areas is certainly part of the reason for this decline, the need for more accurate knowledge of the areas used by wintering migrants for conservation action has been highlighted by numerous authors (Newton, 2008; Morrison et al., 2013; Vickery et al., 2014). Here we show that kestrels spend no less than 40% of the annual cycle in West Africa, and that the average time spent at first, intermediate and last sites equates to 10%, 14% and 20% of the annual cycle, respectively. Our work thus emphasises the conservation value of West Africa for lesser kestrels and of last staging sites in particular, where kestrels aggregate in large numbers close to wetlands. Previous counts in the order of c. 30.000 birds at super-roosts in Central Senegal represented 45% of the population breeding in Western Europe (Pilard et al., 2011; Augiron et al., 2015). Our results indicate that 32% and 49% of lesser kestrels migrating to West Africa from Spain are likely to converge in Senegal and at the Malian-Mauritanian border, respectively. The aggregation of kestrels in large super-roosts at potentially vulnerable habitats in the Sahel (e.g. wetlands, natural savannahs, mosaics) makes the species more sensitive to land use change. Its seeming dependence on insect prey suggests that pesticide use, combined with the continuous conversion of wetlands and grasslands into cropland and excessive forest clearing for firewood by a growing rural population in West Africa, may pose a significant risk to the species (Thiollay 2007, Zwarts et al., 2009).

## Acknowledgements

We would like to acknowledge A. Bermejo and J. de la Puente from SEO/Birdlife for facilitating access to the data. We thank M. Aguilera, E. Aguirre, E. Álvarez, P. Aycart, M. Baena, S. Bondì, F. Carbonell, M.A. Carrero, S. De la Fuente, V. De la Torre, M. Galán, M. Garcés, J.L. González, E. Griffin, L. Hernández, E. Holroyd, D. Jordano, P. Lazo, C. Marfil, J. Marín, F.J. Martín-Barranco, R. Mascara, A. Meijide, P. Moreno, D. Ni Dhubhail, C. Ordóñez, M. Pomarol, F.J. Pulpillo, P. Ruiz, A. Valverde. and L. Zanca. for their help during fieldwork and for technical support. We thank A. Fernández (LIFE project manager in Extremadura) for his support and collaboration. We especially thank M. Vázquez for his support during fieldwork in Doñana.

## Ethics statement

All experimental protocols were approved by Estación Biológica de Doñana Ethical Committee, Consejo Superior de Investigaciones Científicas Ethical Committee, and Consejería de Agricultura, Ganadería, Pesca y Desarrollo Sostenible de la Junta de Andalucía and carried out in accordance with relevant regulations approved by the Spanish Law on Animal Experimentation (RD53/2013 from 1^st^ February https://www.boe.es/eli/es/rd/2013/02/01/53). Capture and device deployment were carried out by experienced ornithologists only in accordance with approved guidelines aimed at ensuring animal welfare throughout the operations (Whitworth et al., 2007).

