## Supplementary methods for "Itinerant lifestyle and congregation of lesser kestrels in West Africa"

### Supporting methods

#### Logger programming

Loggers were programmed with different duty cycles: 50 NanoFix GEO+RF tags had a double schedule. From these, 43 tags collected GPS positions at an interval of 15 min on a 14 h ON/10 h OFF cycle between 6:00 and 20:00 during Jan, May, June, July, Nov, Dec, and at an interval of 30 min on a 24 h ON cycle during Feb, March, April, Aug, Sept, and Oct. Three tags collected GPS positions at an interval of 15 min on a 12 h ON/12 h OFF cycle between 8:00 and 20:00 during May, June, July and at an interval 1 h on a 24 h ON cycle during Jan, Feb, Mar, Apr, Aug, Sep, Oct, Nov, Dec. Four tags collected GPS positions at an interval of 10 min on a 13 h ON/13 h OFF cycle between 8:00 and 21:00 during Mar, Apr, Jun, Jul, Aug, Sep and at an interval of 30 min on a 24 h ON cycle during Jan, Feb, May, Oct, Nov, Dec. Eleven Microsensory tags collected positions at an interval of 15 min on a 15 h ON/9 h OFF cycle between 6:00 and 20:00 and 1 nocturnal position at 1 am. Intervals differed depending on solar battery recharge and satellite geometry (≥ 4 satellites must be detected for a reliable fix).
