## Supplementary figures and images for "Itinerant lifestyle and congregation of lesser kestrels in West Africa"

### Supplemental Figure 1

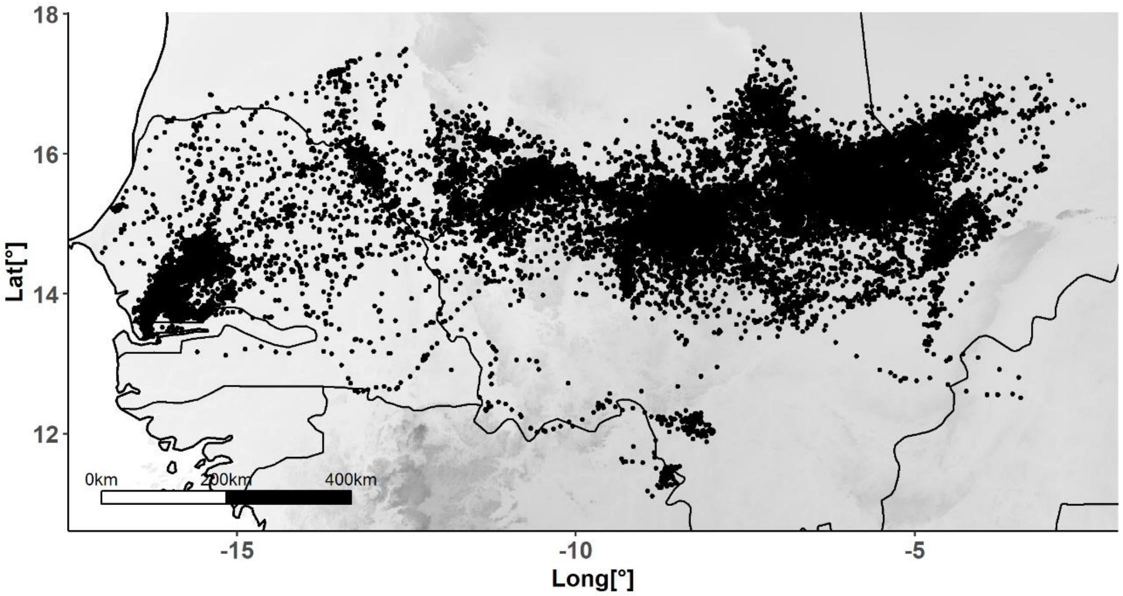


**Supplemental Figure 1.** Map showing all GPS fixes of the 54 lesser kestrels tracked during winter (2016-2020).
