## Supplemental Figure 2 for "Itinerant lifestyle and congregation of lesser kestrels in West Africa"

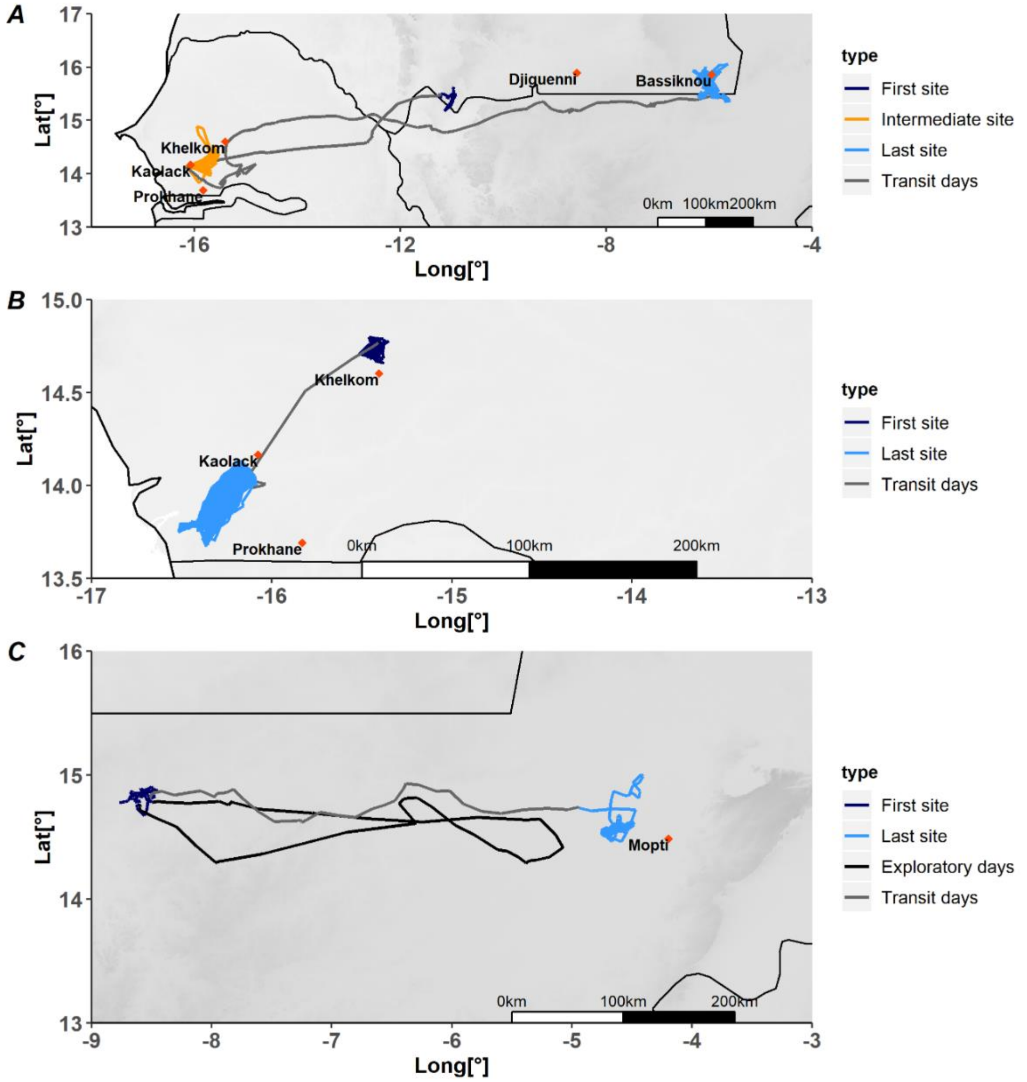


**Supplemental Figure 2.** Example of annotation of GPS tracked adults and movements within the West African wintering area. **(A)** Female lesser kestrel (4170933_R8J4) during the northern winter 2018-2019 used three wintering sites (first site in purple, intermediate in yellow and last in blue). Transit days or days when she flew between sites without returning to the previous site are shown in grey. **(B)** Female (4179802_RU00) arrived at Khelkom (first site) on 2 October 2018. On 27 November, she flew 80 km southward, where she spent most of her winter close to a salt lake near Kaolack, a typical daytime roost known to hold several tens of thousands of raptors (Pilard et al., 2011; Zwarts et al., 2009). She stayed on this wetland until the onset of migration on 17 March 2019. **(C)** Track of a male lesser kestrel (BR.D) during the northern winter (2018-2019) showing exploratory days in black (lasted six days).
