## Supplemental Figure 3 for "Itinerant lifestyle and congregation of lesser kestrels in West Africa"

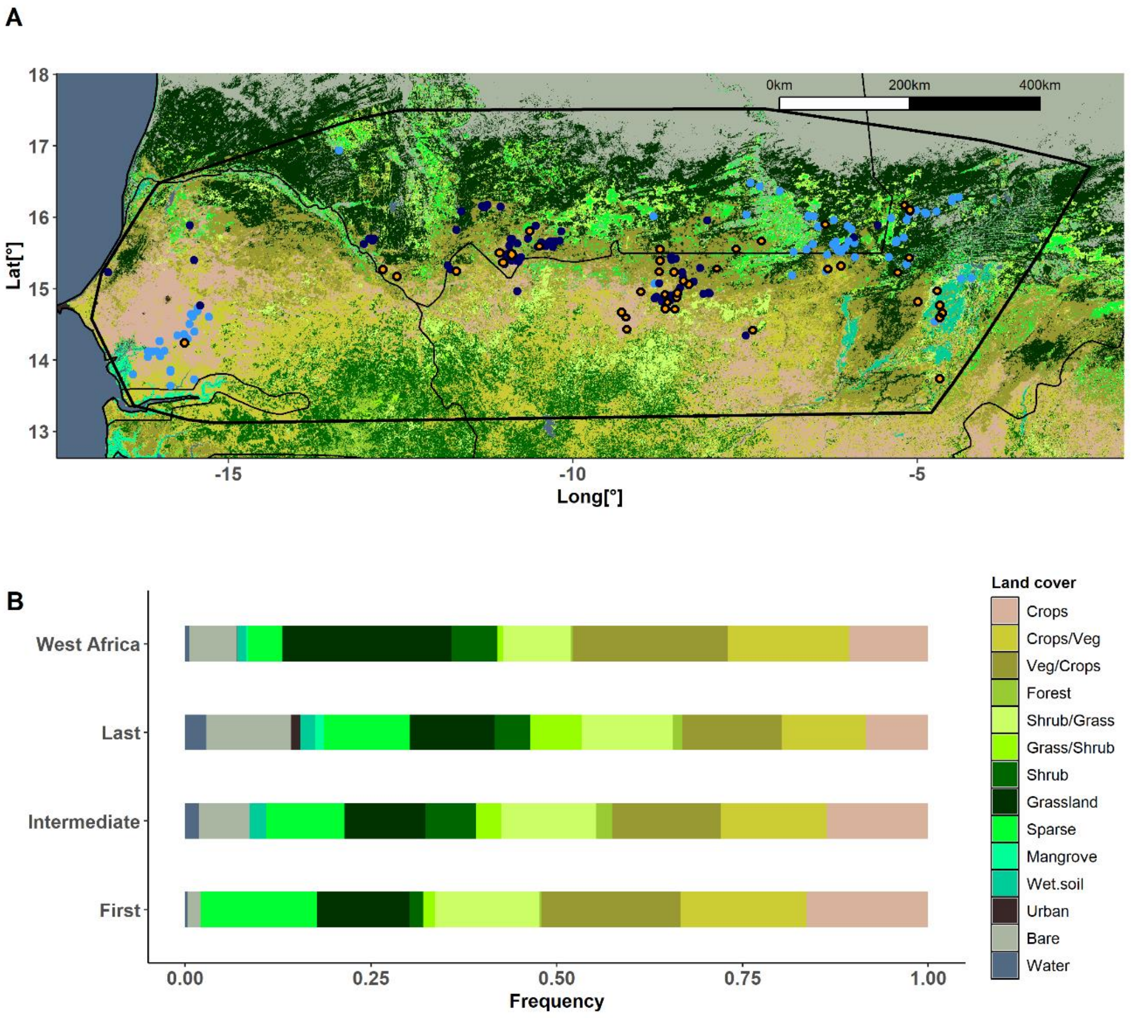


**Supplemental Figure 3. (A)** Habitat composition available in the non-breeding area in West Africa according to the GlobCover land use map. The dots represent the centroids of each staging site: purple dots indicate first sites, orange dots with black border intermediate sites and blue dots last sites. The black polygon indicates 100% MCP for the first, intermediate and last sites. (**B**) Bar graph showing the availability of each land use category in the MCP West African non-breeding range (black-line polygon on map) and the proportion of land cover types used by kestrels (i.e., the proportion of GPS fixes within each land cover category) in the first, intermediate and last staging sites. Fourteen out of 23 GlobCov categories are used by kestrels. Kestrels use crops and mosaics of crops/veg habitats in the first months of their stay in Africa and a shift towards more varied habitat use, including also wetter, urban and bare habitats at last staging sites.
