## Supplemental Table 1 for "Itinerant lifestyle and congregation of lesser kestrels in West Africa"

**Supplemental Table 1.** Wintering summary statistics for 54 lesser kestrels showing: the first year of tracking, sex, numbers of winters tracked, number of staging sites, the average duration of the stay in West Africa, average duration spent at the first, intermediate and last sites, the average number of resident, transit and exploratory days. Average values for birds with 2 or 3 tracks are shown.

| Bird identity | First tracking year | Sex | N of winters tracked | N of staging sites | Mean duration winter period (days) | Mean duration at first sites | Mean duration at intermediate sites | Mean duration at last sites | Resident days | Transit days | Exploratory days |
|---|---|---|---|---|---|---|---|---|---|---|---|
| B00300 | 2018 | f | 2 | 4 | 144 | 27.5 | 46.5 | 63.5 | 130.5 | 6 | 4 |
| B00309 | 2018 | f | 1 | 2 | 141 | 65 | 6 | 66 | 134 | 2 | 2 |
| B00316 | 2018 | f | 1 | 2 | 136 | 18 | | 96 | 81 | 21 | 24 |
| B00323 | 2018 | f | 1 | 3 | 151 | 23 | 107 | 12 | 138 | 13 | |
| B00324 | 2018 | f | 1 | 2 | 144 | 54 | | 80 | 121 | 9 | |
| B00331 | 2018 | f | 1 | 1 | 170 | | | 170 | 102 | | |
| B00336 | 2018 | f | 1 | 3 | 130 | 19 | 43 | 64 | 120 | 3 | |
| B00338 | 2018 | m | 1 | 3 | 152 | 26 | 17 | 31 | 69 | 49 | |
| B00342 | 2018 | m | 2 | 3 | 144 | 35.5 | 34 | 75.5 | 123 | 14 | |
| B00347 | 2018 | f | 1 | 2 | 130 | 3 | 3 | 83 | 63 | 24 | |
| B00348 | 2018 | m | 2 | 3 | 149 | 53.5 | | 68.5 | 97.5 | 12.5 | |
| B00349 | 2018 | m | 2 | 3 | 139 | 23 | | 93 | 115.5 | 18.5 | |
| B16127 | 2017 | m | 2 | 4 | 149 | 26.5 | 104 | 18.5 | 147.5 | 9.5 | 9 |
| B16137 | 2017 | m | 3 | 2 | 140.5 | 64.5 | | 68.5 | 127 | 10 | |
| B16212 | 2016 | m | 1 | 2 | 156 | 61 | | 92 | 148 | 2 | |
| B16228 | 2017 | m | 1 | 2 | 154 | 44 | 20 | 84 | 151 | 4 | |
| B16275 | 2017 | f | 1 | 3 | 175 | 4 | 41 | 113 | 160 | 16 | |
| B16584 | 2017 | f | 1 | 2 | 199 | 54 | 107 | 27 | 191 | 10 | |
| B16611 | 2017 | m | 2 | 2.5 | 177 | 62.5 | | 33 | 170.5 | 6 | |
| B16639 | 2017 | m | 1 | 2 | 149 | 66 | | 81 | 148 | 1 | |
| B16643 | 2017 | f | 2 | 2 | 132.5 | 45 | | 85.5 | 131 | 1 | 3 |
| B16645 | 2017 | f | 2 | 3 | 138 | 19.5 | 78.5 | 27.5 | 121 | 16 | |
| B16661 | 2017 | f | 2 | 1.5 | 158 | 65 | | 124.5 | 140.5 | 1 | 2 |
| B16679 | 2017 | f | 2 | 2 | 171.5 | 21.5 | 36 | 99 | 154 | 14 | 2 |
| B16687 | 2017 | f | 1 | 2 | 154 | 3 | | 149 | 154 | 1 | |
| B16688 | 2017 | m | 2 | 2.5 | 139 | 66.5 | | 70 | 137 | 1.5 | 3 |
| B16690 | 2017 | m | 2 | 3.5 | 165.5 | 24 | 50 | 81.5 | 154.5 | 10.5 | 3 |
| B17199 | 2018 | f | 1 | 3 | 126 | 3 | 77 | 40 | 121 | 6 | |
| B17210 | 2018 | m | 2 | 2 | 105 | 19.5 | 30 | 61.5 | 95.5 | 7.5 | 6 |
| B17214 | 2018 | f | 1 | 4 | 139 | 17 | 94 | 24 | 133 | 8 | |
| B17215 | 2018 | f | 1 | 2 | 147 | 11 | 26 | 89 | 129 | 19 | |
| B17218 | 2018 | m | 1 | 4 | 135 | 5 | 106 | 20 | 121 | 15 | |
| B17219 | 2018 | m | 1 | 2 | 142 | 5 | | 135 | 142 | 8 | |
| B17230 | 2018 | m | 2 | 2.5 | 108 | 10 | 17 | 77 | 94.5 | 13.5 | 2 |
| B17235 | 2018 | m | 2 | 2 | 143.5 | 63.5 | | 73 | 135.5 | 6 | 6 |
| B17237 | 2018 | m | 2 | 3.5 | 153 | 35 | 102.5 | 8.5 | 141 | 13.5 | |
| B17239 | 2018 | f | 2 | 2 | 146 | 11.5 | 22.5 | 91.5 | 128.5 | 16.5 | 7 |
| B17240 | 2018 | m | 1 | 2 | 151 | 51 | | 90 | 129 | 9 | 14 |
| B17241 | 2018 | f | 1 | 4 | 171 | 65 | 92 | 9 | 164 | 8 | |
| B17245 | 2018 | f | 1 | 2 | 132 | 66 | | 24 | 92 | 41 | |
| B17250 | 2018 | m | 2 | 2.5 | 146.5 | 42.5 | 62 | 67 | 141 | 5.5 | |
| B17251 | 2018 | m | 1 | 3 | 129 | 26 | 52 | 47 | 127 | 3 | |
| B17253 | 2018 | m | 2 | 1.5 | 145.5 | 67 | 14 | 93.5 | 130 | 12.5 | |
| B17254 | 2018 | m | 2 | 3 | 146 | 17 | 88 | 35 | 140 | 9 | 1 |
| B17256 | 2018 | f | 1 | 2 | 114 | 31 | 31 | 39 | 104 | 11 | |
| B17261 | 2018 | m | 1 | 2 | 156 | 48 | 38 | 19 | 108 | 49 | |
| B17262 | 2018 | f | 1 | 3 | 151 | 49 | 50 | 46 | 134 | 18 | |
| B17278 | 2018 | f | 2 | 2.5 | 160 | 27.5 | 51 | 103 | 155 | 5.5 | |
| B17280 | 2018 | f | 1 | 2 | 154 | 39 | 58 | 41 | 141 | 14 | |
| B17281 | 2018 | f | 1 | 2 | 150 | 24 | 42 | 80 | 149 | 2 | |
| B17282 | 2018 | f | 2 | 3 | 136.5 | 29 | 62.5 | 41 | 128 | 4 | 11 |
| B17288 | 2018 | f | 1 | 2 | 166 | 72 | 33 | 54 | 161 | 10 | 1 |
| B17292 | 2018 | f | 1 | 2 | 165 | 55 | | 108 | 165 | 1 | |
| B40128 | 2018 | m | 2 | 2 | 134 | 40 | 20 | 80 | 132.5 | 2.5 | |
