## Supplemental Table 2 for "Itinerant lifestyle and congregation of lesser kestrels in West Africa"

**Supplemental Table 2. GlobCover legend description**

| GlobCover value | GlobCover label | Description | Abreviation |
|---|---|---|---|
| 14 | Rainfed croplands | Rainfed shrub crops/ rainfed tree crops / rainfed herbaceous crops | Crops |
| 20 | Mosaic cropland: 50-70% cropland / 20-50% vegetation (grassland, shrubland, forest) | Cultivated and managed terrestrial areas / natural and semi-natural, primarily terrestrial vegetation | Crops/Veg |
| 30 | Mosaic vegetation: 50-70% vegetation (grassland, shrubland, forest) / 20-50% cropland | Natural and semi-natural, primarily terrestrial vegetation / cultivated and managed terrestrial areas | Veg/Crops |
| 40 | >15% closed to open broadleaved evergreen and/or > 5m semi-deciduous forest | Broadleaved evergreen closed to open trees / semi-deciduous closed to open trees | Forest |
| 60 | 15-40% open broadleaved deciduous forest/ >5m woodland | Broadleaved deciduous (40-(20-10)%) woodland | |
| 110 | Mosaic shrubland: 50-70% shrubland / 20-50% grassland | Closed to open trees / closed to open shrubland (thicket) / herbaceous closed to open vegetation | Shrub/Grass |
| 120 | Mosaic grassland: 50-70% grassland / 20-50% forest or shrubland | Closed to open shrubland (thicket) / herbaceous closed to open vegetation / closed to open trees | Grass/Shrub |
| 130 | >15% closed to open (broadleaved or needleleaved, evergreen or deciduous) <5m shrubland | Broadleaved closed to open shrubland (thicket) | Shrub |
| 140 | >15% closed to open herbaceous vegetation (grassland, savannas or lichens/mosses) | Herbaceous closed to very open vegetation / Closed to open lichens/mosses | Grassland |
| 150 | <15% sparse vegetation | Sparse trees / herbaceous sparse vegetation / sparse shrubs | Sparse |
| 170 | >40% closed broadleaved forest or shrubland permanently flooded - saline or brackish water | Closed to open (100-40%) broadleaved trees on permanently flooded land (with daily variations), water quality: saline water / closed to open (100-40%) broadleaved trees on permanently flooded land (with daily variations), water quality: brackish water / closed to open (100-40%) semi-deciduous shrubland on permanently flooded land (with daily variations), water quality: saline water / closed to open (100-40%) semi-deciduous shrubland on permanently flooded land (with daily variations), water quality: brackish water | Mangrove |
| 180 | >15% closed to open grassland or woody vegetation on regularly | Closed to open shrubs / closed to open herbaceous vegetation | Wet.soil |

| | | | |
|---|---|---|---|
| | flooded or waterlogged soil - fresh, brackish or saline water | | |
| 190 | >50% urban areas | Artificial surfaces and associated areas | Urban |
| 200 | Bare areas | Bare areas | Bare |
| 210 | Water bodies | Natural water bodies / artificial water bodies | Water |
